# *In vitro* models of the human esophagus reveal ancestrally diverse response to injury

**DOI:** 10.1101/2021.05.20.444920

**Authors:** Daysha Ferrer-Torres, Joshua H. Wu, Charles J. Zhang, Max A. Hammer, Michael Dame, Angeline Wu, Emily M. Holloway, Kateryna Karpoff, Caroline L. McCarthy, Margaret S Bohm, Sha Huang, Yu-Hwai Tsai, Simon P. Hogan, Danielle Kim Turgeon, Jules Lin, Peter D.R. Higgins, Jonathan Sexton, Jason R. Spence

**Affiliations:** Department of Internal Medicine, Division of Gastroenterology; University of Michigan Medical School, Ann Arbor, Michigan; Department of Cell and Developmental Biology; University of Michigan Medical School, Ann Arbor, Michigan; Department of Biomedical Engineering; University of Michigan Medical School, Ann Arbor, Michigan; Center for Cell Plasticity and Organ Design, University of Michigan Medical School, Ann Arbor, Michigan; Department of Medicinal Chemistry, College of Pharmacy, University of Michigan, Ann Arbor, Michigan; Department of Thoracic Surgery, University of Michigan Medical School, Ann Arbor, Michigan; Department of Pathology, University of Michigan Medical School, Ann Arbor, Michigan

**Keywords:** esophagus, adult patient-derived cultures, biobank, disparity, ancestry

## Abstract

European Americans (EA) are more susceptible to esophageal tissue damage and inflammation when exposed to gastric acid and bile acid reflux and have a higher incidence of esophageal adenocarcinoma when compared to African Americans (AA). Population studies have implicated specific genes for these differences; however, the underlying cause for these differences is not well understood. We describe a robust long-term culture system to grow primary human esophagus *in vitro*, use single cell RNA sequencing to compare primary human biopsies to their *in vitro* counterparts, identify known and new molecular markers of basal cell types, and demonstrate that *in vivo* cellular heterogeneity is maintained *in vitro*. We further developed an ancestrally diverse biobank and a high-content, image based, screening assay to interrogate bile-acid injury response. These results demonstrated that AA esophageal cells responded significantly differently than EA-derived cells, mirroring clinical findings, having important implications for addressing disparities in early drug development pipelines.

## Introduction

The esophagus connects the upper pharynx with the stomach and is lined by a stratified squamous epithelium (Rosekrans *et al*., 2015). The esophagus is prone to many diseases including esophageal squamous cancer (Kim *et al*., 2017), eosinophilic esophagitis (Blevins *et al*., 2018), metaplasia (Barrett’s esophagus) due to reflux and inflammation, and esophageal adenocarcinoma (Saraggi *et al*., 2016). Despite risk factors being equal across populations (Spechler *et al*., 2002; El-Serag *et al*., 2004), there is a higher incidence of esophageal adenocarcinoma in the Caucasian/European American (EA) population when compared to populations of African descent, including the Black/African American (AA) population (El-Serag, HB and Sonnenberg, 1997; Rastogi *et al*., 2008; Sharma *et al*., 2008; El-Serag *et al*., 2014; Arnold *et al*., 2017; Then *et al*., 2020). Genetic variations attributed to ancestry and racial background has also been implicated in various genetic diseases (i.e. sickle cell anemia (Kwiatkowski, 2005), cystic fibrosis (Knowles and Drumm, 2012), risk of EAC development (Ferrer-Torres *et al*., 2019) and may lead to differences in response to therapeutic drugs (Ortega and Meyers, 2014; Hunter, 2020). Therefore, inclusion of individuals from diverse demographics for drug development and clinical trials is important for ensuring efficacy across the population. Although mandated by federal regulations and NIH policy, the inclusion of a diverse population is still aspirational, and appropriately addressing racial disparities has faced criticism as recently as the current COVID19 pandemic. Thus, failing to study racially/ancestrally and ethnically diverse populations will leave critical gaps in our understanding of variation to drug responses, and the effectiveness of new therapies. In the context of esophageal disease, genetic studies suggest racial differences in the tissue response to cell and DNA damage-inducing agents leading to carcinogenesis (Ferrer-Torres *et al*., 2019); however, the ability to study racial disparities in esophageal disease has been hindered by the lack of diverse human models to study these mechanisms.

In this study we aimed to first characterize the heterogenous cell types found in the healthy/normal human esophagus *in vivo*, to establish a racially/ancestrally diverse esophageal biobank of *in vitro* primary tissue lines, and to compare *in vivo* and *in vitro-*derived cultures using single-cell RNA sequencing (scRNA-seq). We developed a diverse biobank including tissue from individuals self-identified as EA, AA, Asian, and Hispanic descent (n=55 cell lines)(Table 1). Using scRNA-seq, we identified four molecularly distinct zones within the native *in vivo* esophageal epithelium and validated new and known markers for each zone: basal (COL17A1^+^, CAV1^+^, CAV2^+^), suprabasal (LY6D^+^), mid-suprabasal (KRT4^+^), luminal zone (CRNN^+^). We found that *in vitro* cultured cells could be propagated for many generations, and while these cultures possessed an abundance of basal stem cells, they also recapitulated the cellular diversity observed in the *in vivo* esophagus, including early differentiating squamous cells. Secondly, we aimed to understand how these heterogenous cell types respond to bile-acids. Multiple studies in animals have shown that bile acids, but not stomach acid alone, contribute to the formation of columnar and glandular tissue, characteristic of Barrett’s Esophagus (Kauer *et al*., 1995; Quante *et al*., 2012; Straub *et al*., 2019). Therefore, we developed an image-based 384-well high content screening assay and image analysis pipeline yielding single-cell phenotypic measurements. Machine learning was then used to interrogate cellular phenotypes and showed that AA cells are less sensitive to damage by exposure of bile-acid, mimicking bile-acid reflux. These results are consistent with clinical observations and suggest that *in vitro* human model systems can capture genetic diversity leading to different biological response to injury

**Table 1.**
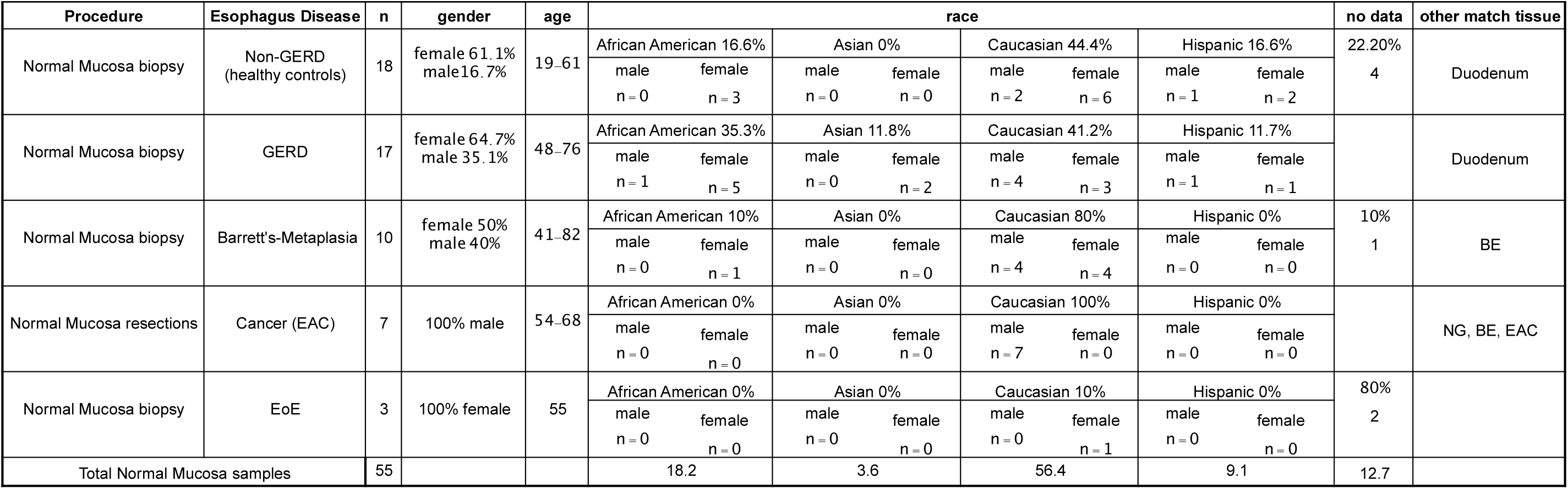
Clinical Characteristics of Primary Human Esophageal Samples for 2D or 3D *in vitro* Culture.

## Results

### Human adult esophagus epithelium contains molecularly-distinct zones defined by scRNAseq and validated at the protein level

The esophagus its composed of a stratified squamous epithelium, underlying stromal tissue (lamina propria) and smooth muscle (muscularis mucosae)(Rosekrans *et al*., 2015). In order to characterize the esophagus, we obtained normal/healthy squamous epithelial (normal squamous - “NS”) adult human biopsies (approximately 3mm^2^) (n=2 independent patients, with n=3-4 biopsies used for dissociation), carried out enzymatic dissociation into single cells, which were captured using the 10X Chromium platform for subsequent sequencing. Louvain clustering analysis defined seven transcriptionally distinct clusters (Figure S1A-C, Table S1), and defined classes of cells as Epithelial (*CDH1*^+^ - Clusters 0,1,2,3) or stroma/lamina propria (*VIM*^*+*^ - Clusters 4,5,6) which included immune cells (Clusters 4/5) (Figure S1B-F). Contribution to each cluster was consistent across both biological replicates (Figure S1B). The top 5 genes for each cluster were plotted (Figure S1E), and other enriched genes for each cluster, which included known genes (Table S1) were used to identify cells associated with different zones of the esophageal epithelium (Figure S1E-F). Cluster 3 is characterized by markers expressed within the basal zone of esophagus (Figure S1E-F), and Cluster 1 shares expression of basal cell markers but also includes a proliferative signature (Figure S1E-F). We also identified KRT4-positive clusters (Clusters 0, 2) cells which mark the transitional and luminal populations of the squamous epithelium. Cluster 2 expressed CRNN and CNFN, which are indicative of the most differentiated cells at luminal surface of the epithelium. Many genes identified in scRNA-seq clusters were further screened at the protein level using the Human Protein Atlas (HPA) (Uhlén *et al*., 2005, 2015), demonstrating that molecular identities correlated with different zones determined by protein staining for different layers (basal, suprabasal, luminal) within the stratified epithelium (Figure S1G).

To further characterize the epithelium, *CDH1*-positive clusters were computationally extracted and reclustered, revealing four predicted sub-clusters (Figure 1A, Table S2). Individual biopsies contributed to each cluster in similar proportions (Figure 1B). Unsupervised clustering was used to plot the top 5 genes in an unbiased way (Figure 1C, Table S2) and used to determine that epithelial clusters correspond to basal cells (Cluster 2), proliferative (Cluster 1), suprabasal (Cluster 0), and a differentiated/luminal zone cells (Cluster 3) (Figure 1C, Figure S2A). Of note, we observed that *TP63*, a marker canonically used to identify basal cells, was broadly expressed in basal, proliferative and transitional cell clusters (Figure S2B), a finding that was validated with protein staining, which showed broad epithelial expression (Figure S2C). scRNA-seq data identified genes that were highly enriched in the basal cell cluster (Cluster 2)(Figure 1C-D), which had specific localization to the basal cell layer by immunofluescence, and included CAV1, CAV2, and COL17A1 (Figure 1C-E, Figure S2). scRNA-seq identified *LY6D* as an enriched marker in the suprabasal/early transitional squamous and proliferative cluster (Clusters 0, 1) (Figure 1D, Figure S2). Supporting this, immunofluorescence shows that LY6D protein is localized to the suprabasal zone just above the basal cell domain (Figure 1E, Figure S2C), which is also where the majority of KI67+ proliferative cells are observed (Figure 1E). Within the suprabasal cluster (Cluster 0), we observed that *KRT4* is expressed in a low-to-high gradient (Figure 1D), and at the protein level, KRT4 marks the mid-point of the transitional zone, above the LY6D+ epithelium (Figure 1E, Figure S2C). Finally, Cluster 3 represents a CRNN-high zone that marks differentiated cells in the portion of the squamous epithelium near the lumen (Figure 1C-E; Figure S2C). Altogether, we have mapped the epithelial zones of the esophagus at the mRNA and protein level, and identified markers for each zone, with CAV1, CAV2 and COL17A1, uniquely marking the basal cells.

**Figure 1.**
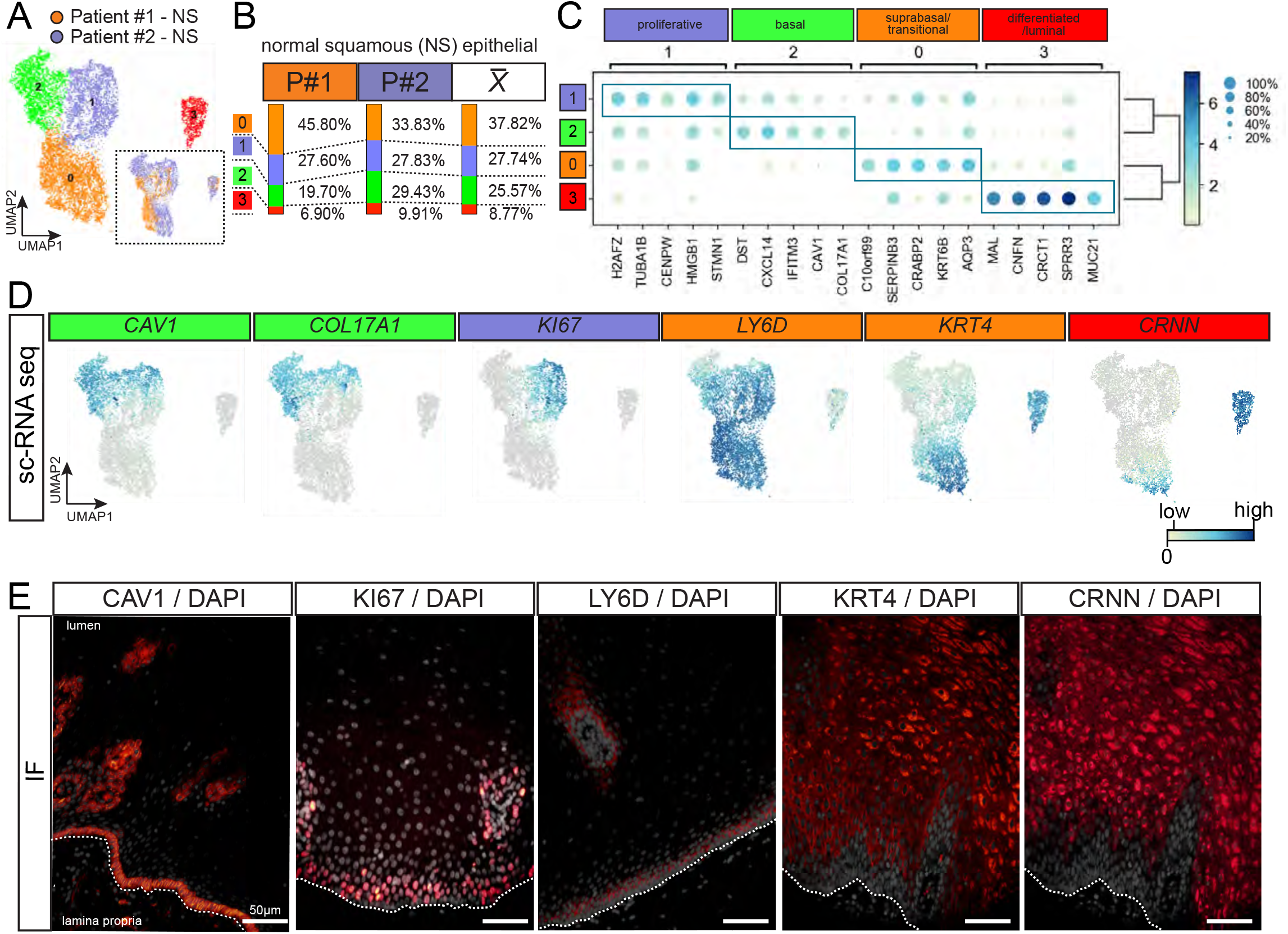
Identification of distinct molecular domains within the esophageal epithelium. (A) *CDH1+* epithelial cells were sub-clustered from scRNA-seq data for patient esophageal biopsies (n=2) with a total of 7796 cells were analyzed after filtering, and 2651 genes per cell. Louvain clustering was used to predict clusters and visualized using UMAP. (B) Distribution of patient cells to each cluster. <u>*x*</u> denotes the average contribution of both samples to the clusters. (C) Dot-plots of the top 5 genes expressed in each cluster, and annotations for each cluster based on top genes and known genes (see also Table S2). (D) Feature plots of top marker genes expressed in each cluster, with *CAV1/COL17A1* for Cluster 2 (basal), *KI67* for Cluster 1 (proliferative), *LY6D* and *KRT4* for Cluster 0 (suprabasal/transitional), and *CRNN* for Cluster 3 (differentiated/luminal). (E) Representative immunofluorescent images in the adult human esophagus validating genes identified by scRNA-seq. The markers localize COL17A1, CAV1, CAV2 (see Figure S2) at the basal zone, KI67 marks proliferative cells at the basal-suprabasal zone, LY6D is negative in the basal layer and marks the first suprabasal layer, KRT4 is an early differentiation marker, and CRNN stains the luminal/cell layer of fully differentiated cell types. (Other markers validated in Figure S2). Scale bars represent 50 ⌈m (Images are representative of n=3 biological replicates).

### Long term maintenance of patient-derived human esophageal basal-stem cells in vitro

In order to create a robust biorepository of diverse esophagus epithelial cell lines, we modified previously established methods for deriving *in vitro* epithelial cell cultures that included sub-lethally irradiated 3T3-J2 feeder cells (X. Liu *et al*., 2017) coupled with “dual SMAD inhibition” media (Mou *et al*., 2016). On the day of biopsy (Day 0 – D0), tissues were finely minced and plated on the irradiated feeder (3T3-J2i) layer with media (see Methods). Cell clumps attached and expanded as small colonies, which eventually grew to confluence (Figure 2A-B). We observed that the irradiated 3T3-J2i cells do not proliferate, while esophageal cells continue proliferating over time (Figure 2C). After three passages, and 30-40 days in culture, we assessed cultures of these patient-derived *in vitro* esophagus cells using scRNA-seq (n=3). We applied Louvain clustering to reveal five molecular clusters (Figure 2D, Table S3), and plotted the proportion of total cells within each cluster (Figure 2E). All clusters were enriched for the epithelial marker *CDH1* expressed (average of biological replicates *CDH1+* 96.6% vs *VIM+* 5.4%, *(Figure S3E*)) and protein staining revealed a high percentage of cells within the culture were ECAD+ (Figure 1F-G), a finding that was quantitated (% ECAD+ cells, n=7 independent lines, Figure 2H top) demonstrating ∼50-80% of cells were ECAD+, and that this did not change across passage (Figure 2H, bottom). The top 5 most highly enriched genes were plotted (Figure S3A) and identified that Cluster 4 contains a proliferative signature, Clusters 1 and 2 expressed markers of basal-stem cells including *CAV1, CAV2* and *COL17A1* and known markers such as *TP63, KRT15* and *ITGB4* (Figure 2I, Figure S3B-C) while Clusters 0 and 3 expressed markers of the suprabasal and luminal zone (Figure 2I). *ITGB4* and *COL17A1* expression is consistent with previous work showing that it is a stem-cell marker of esophagus cells (DeWard, Cramer and Lagasse, 2014; Bogte *et al*., 2021). We validated protein expression of basal cell markers, COL17A1, CAV1, and CAV2 and demonstrated that these cells continue to co-express TP63 *in vitro* (Figure 2J, Figure S3 D, F-G). Finally, we quantified TP63^+^/KI67^+^ cells culture, and observed that low density cultures grew as highly proliferative colonies (n=3), while high density cultures formed a monolayer with significantly reduced proliferation (Figure 3K), an observation that was validated in three independents cell lines. Together, these data suggest that human esophagus epithelial cells are maintained in long-term culture, and that they retain many molecular markers seen *in vivo*.

**Figure 2.**
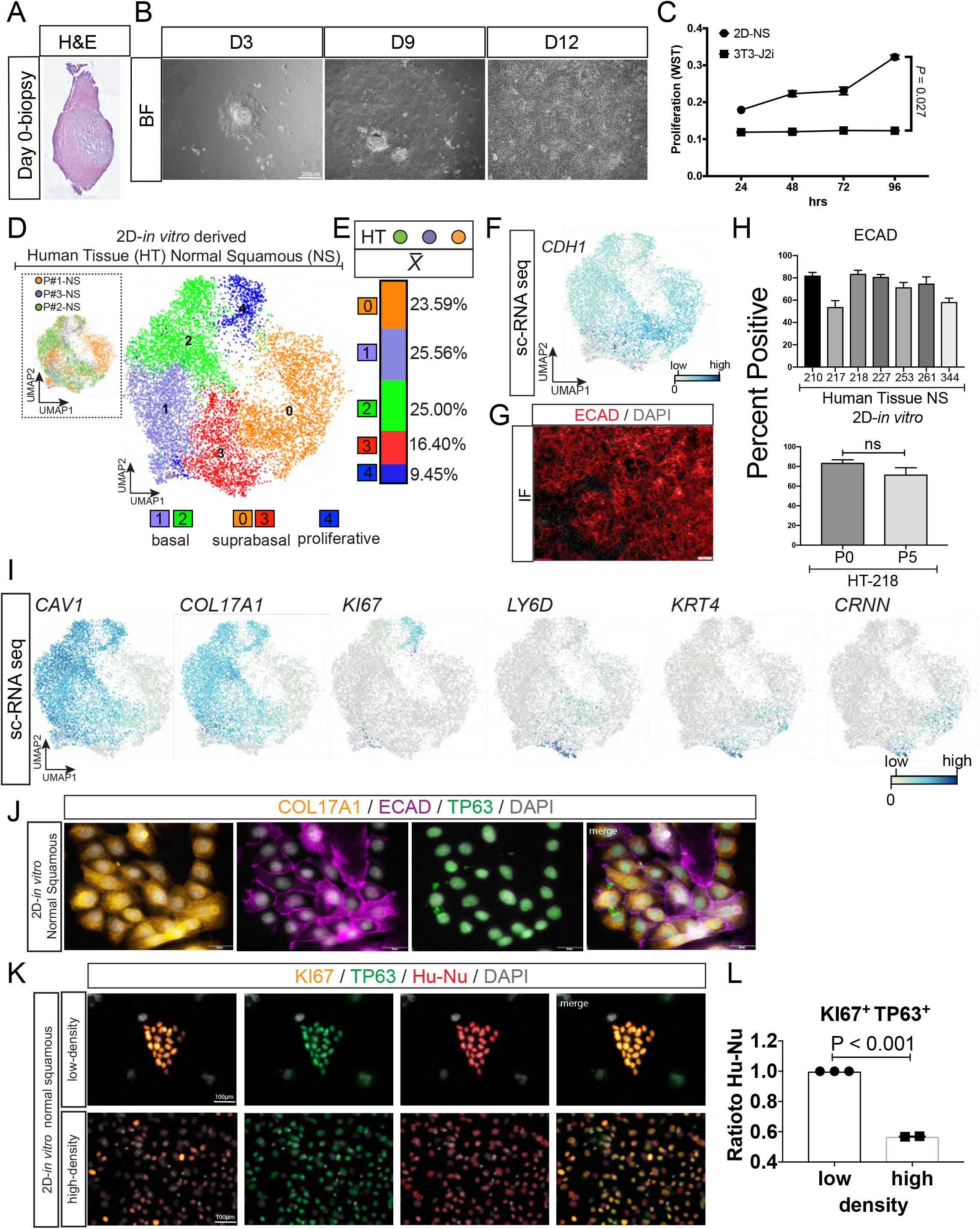
Characterizing human esophageal biopsies grown *in vitro*. (A) H&E of a representative biopsy of squamous epithelial cells of the esophagus. (B) Bright field (BF) images of expansion of esophagus cell clusters/colonies for Day 3, Day 9 and Day 12. Scale bar represents 200 µm. (C) Proliferation as measured using the WST assay at 24, 48, 72, and 96hrs where esophageal cells are proliferating over time compared to sub-lethally irradiated 3T3-J2 mouse fibroblast cells (t-test *P*=0.027). (D) A total of 10550 cells grown *in vitro* and 4269 genes/cell were analyzed using Louvain clustering and visualized via UMAP to predict five clusters. C1/C2 express basal cell markers (See Figure 2I), C0/3 express markers of early suprabasal cells, and C4 express proliferation markers. (Table S3) (E) Distribution plot of the average (<u>*x*</u>) number of cells contributing to each cluster per sample (n=3 biological replicates). (F-H) Characterization of the epithelium in *in vitro* cultures. (F) *CDH1* expression across clusters, with (G) validation of expression of ECAD (protein) suggesting an enrichment of epithelial cell types using these methods. (H) Quantification of the % of total cells that are ECAD+ *in vitro*, in multiple patient-derived cell lines (each number on x-axis represents a unique patient sample) (top panel). Epithelial cells do not significantly increase or decrease over passage number (bottom panel) (n=3 t-test, *P* = not significant). (I) Feature plots of genes expressed in basal (*CAV1/COL17A1*), proliferative (*KI67*), suprabasal (*KRT4*), and luminal cells (*CRNN*) (J) Co-expression of the basal cell marker COL17A1 with ECAD and TP63 *in vitro*. 50 µm (K) At low density, Human nuclei are identified by human specific nuclear antigen (Hu-Nu; red), and co-stained with TP63 (green) are highly proliferative, marked by KI67, when compared to the same cell line plated at higher density (bottom row). Scale bars 100µm. (L) Quantification of KI67+/TP63+ cells in low and high confluence. Less confluent cell colonies are highly proliferative compared to high density, confluent monolayers (n=3; t-test; *P* < 0.001). All experiments were performed using at least n=3 biological replicates and t-test was used to compare the mean of groups.

**Figure 3.**
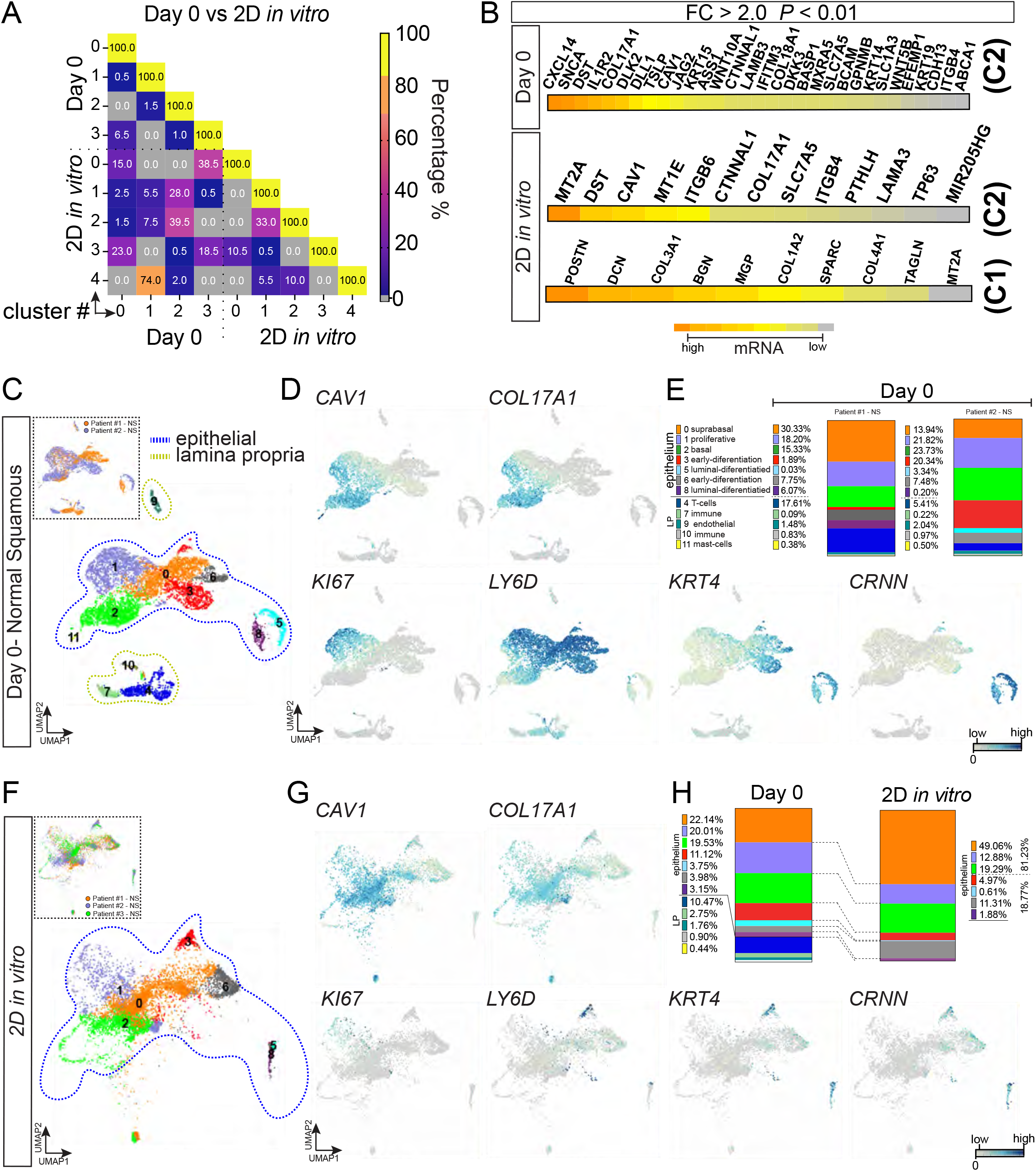
*In vitro* and *in vivo* esophagus share a high degree of molecular similarity. (A) Heat-map representing a direct comparison of the top 200 genes in each cluster for *in vivo* Table S2) and *in vitro* (Table S3) esophageal cells. The % of genes overlapping between lists is plotted, grey boxes have zero overlap. (B) Heat map showing the top genes (FC>2, P<0.01) for *in vivo* basal cells (Figure 1A, Cluster 2 – C2) and for *in vitro* basal cells (Figure 2D - Cluster 1 (C1) and Cluster 2 (C2)). *DST, CAV1, COL17A1, ITGB4, CTNNA1L* are represented in both groups. (C) Louvain clustering and UMAP visualization of *in vivo* samples (blue-dotted line highlighting the epithelial (*CDH1+*) cluster vs. yellow-dotted line highlighting *VIM+* cells. with (D) Feature plots of genes expressed in different cells within the esophagus, including *CAV1, COL17A1 KI67, LY6D, KRT4* and *CRNN*. (E) Distribution of cells from each human sample to each cluster. (F) The Scanpy function Ingest was used to project *in vitro* grown cells onto the *in vivo* cell embedding. *In vitro* cells map to 5 clusters, with the majority of cells mapping to *in vivo* basal (Cluster 2) and suprabasal clusters (Cluster 0). (G) Feature plots showing expression of basal cell genes (CAV1, COL17A1), proliferation associated genes (KI67), suprabasal marker genes (LY6D) and differentiated marker genes (KRT4, CRNN). (H) Quantification of the proportion of cells from the *in vivo* sample in each cluster and the proportion of *in vitro* cultured cells that map to each *in vivo* cluster, demonstrating that *in vitro* cultured cells maintain similar basal cell proportions (Cluster 2, green) but have a larger proportion of suprabasal-like cells (Cluster 0, orange).

### Human-derived esophageal basal-stem cells molecularly resemble native tissue

To further interrogate how closely *in vitro* samples resemble *in vivo* esophageal tissue, we carried out three complimentary but separate analyses using the data. First, we directly compared the single cell transcriptomes of *in vivo* (Figure 1) and *in vitro* (Figure 2) samples (Figure 3A-B); secondly, we integrated *in vitro* and *in vivo* data following batch correction to analyze the samples in one analysis (Figure S4A-E); third, we performed label transfer using Ingest (Wolf, Angerer and Theis, 2017), using the *in vivo* UMAP embedding as a high dimensional search space and projecting *in vitro* samples onto the *in vivo* map (Figure 3C-H). We directly compared the clusters from Day 0 (fresh biopsies) vs 2D *in vitro* cultures (Figure 3A-B) and observed significant overlap of the basal enriched genes observed *in vitro* compared to the *in vivo* cluster. More specifically, we observed an enrichment of the same molecular signature in the proliferative clusters (74% overlap) and the basal clusters (not that there were two basal clusters *in vitro* vs. one cluster *in vivo*, with 39% and 33% overlap, respectively) (Figure 3A-B). Finally, the suprabasal cluster and differentiated clusters *in vivo* (Clusters 0 and 3) had the highest shared similarity to the differentiating *in vitro* clusters (Clusters 0 and 3)(Figure 3A). With respect to the basal cell clusters (C2 *in vivo* vs C1 and C2 *in vitro*), overlap included well established basal cell markers that were common between clusters (i.e. *CAV1, CAV2, ITGB4, COL17A1*) (Figure 3B). Next, we directly compared samples by integrating and batch correcting *in vivo* data (epithelium only) and *in vitro* data with BBKNN (Polański *et al*., 2020), followed by clustering (Figure S4A-E). Integrated data generated four predicted clusters (Figure S4B-C), which could be assigned to proliferative (Cluster 3), basal (Cluster 0), suprabasal (Cluster 1) and luminal (Cluster 2) cell types based on expressed genes (Figure S4B-F). Both *in vitro* and *in vivo* cells contributed to each cluster (Figure S3A), however; the distribution from *in vitro* or *in vivo* cells was not equal for all clusters. For example, only ∼4.5% of cells were designated as basal cells (Cluster 1) from the *in vivo* sample whereas ∼37.4% of cells were assigned to this cluster from the *in vitro* sample (Figure S1E). This observation is consistent with our individual analysis of *in vitro* scRNA-seq data and confirmatory immunofluorescence (Figure 2) showing an abundance of basal cells in these cultures. Lastly, we re-clustered the *in vivo* data (entire data set, including stroma/immune), and assigned identities to the clusters (Figure 3C-E), and then used Ingest to map the location of the *in vitro* cells onto the *in vivo* map (Figure 3F-H). This analysis revealed that the majority of *in vitro* cells mapped to the proliferative, basal and transitional suprabasal cell clusters of the *in vivo* search space, with far fewer cells mapping to the differentiated luminal cells (Figure 3G-H), and as highlighted by distribution plots comparing the proportion of cells from *in vivo* tissue to each cluster versus the projected proportion from *in vitro* cells (Figure 3H).

### A high content bile-acid injury assay models racial disparities in EA and AA esophageal cell lines

High bile-acid content in gastric reflux has been associated with development and a higher incidence of metaplastic progression (Stein *et al*., 1994; Nehra *et al*., 1999). This suggests damage to the normal squamous esophageal mucosa, which precedes the development of metaplasia, is an important target for this bile-acid induced damage. The cell of origin of BE is still highly debated in the field (Que *et al*., 2019) but the role of bile acids is well established as a significant contributor to the development of BE and consequently, EAC. Since the first tissue that is exposed to the reflux is the initially healthy esophageal squamous epithelium, we wanted to test if AA-derived esophageal cells respond differently to bile acid injury than EA-derived cells.

To this end, we developed a bile acid injury assay compatible with high-content imaging based screening, coupled with automated image analysis of cellular features using a Cell Painting (Bray *et al*., 2016) style approach, followed by cell-level machine learning to characterize the perturbation of cells to a mix of primary and secondary bile-acids known to be present in humans (bile-acid mix (BAM))(Nehra *et al*., 1999; Straub *et al*., 2019) see methods) (Figure 4A, Figure S5). First, we tested the individual and mixed bile-acids identified in human reflux (Straub *et al*., 2019) versus oxidative damage induced by Cumene hydroperoxide which has been shown to cause oxidative stress induced DNA damage in the esophagus (Peng *et al*., 2014, 2021; D. Ferrer-Torres *et al*., 2019). We observed that after 48hrs BAM has similar effects on cells when compared to Cumene hydroperoxide, and when compared to individual bile acids (Figure S5A-B). Therefore, we proceeded to test the response of EA and AA cells at 48hrs with increasing concentrations of BAM. After 48hrs of exposure to BAM, a visible morphological change occurred in all patient-derived cell lines (Figure 4B and Figure S5B-C) and we observed that cells have reduced area, are elongated and exhibit increased branching (Figure 4B and Figure S5B-C). Following exposure, cells were fixed and stained with Hoechst 33342, Cell Mask Orange, and for a pro-inflammatory marker NFkB (Jenkins *et al*., 2004; Huo *et al*., 2017; T. Liu *et al*., 2017) (Figure 4B). Automated image analysis was performed using CellProfiler (McQuin *et al*., 2018) to obtain ∼1,400 morphological features of each cell.

**Figure 4.**
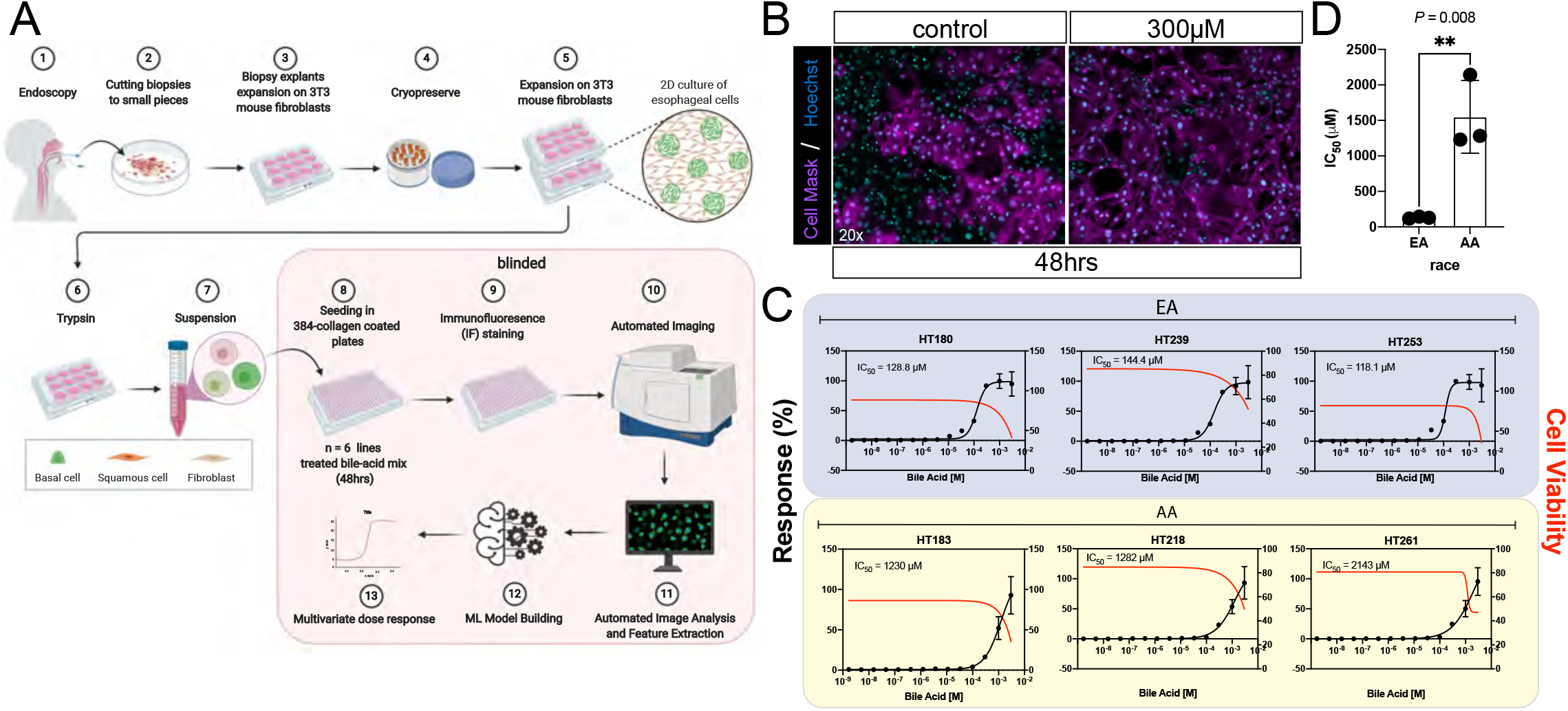
Cells derived from EA versus AA patients respond differently to Bile-acid mix (BAM)-induced injury. (A) Schematic of protocol for esophagus cell expansion and high-content screen. Studies are blinded, and Researcher 1 carried out all cell culture work. Suspensions of coded cells are given to Researcher 2, who performed bile-acid dose response, staining, imaging and data analysis. (B) Representative immunofluorescence images of control (DMSO) vs. 3000 μM bile-acid mix treated cells demonstrating morphological changes from bile-acid exposure (48hrs). (C) High-content imaging on representative cell lines treated through a dose range of bile acid mixture (0 to 3000 μM) followed by automated image analysis for the extraction of 1,400 morphological features per-cell level. Ensemble learning through random forest model was used to classify predicted exposure level of bile-acid per-cell as a percent response of 3000 μM bile-acid mix treated cells. Data shown is average predicted exposure vs actual treatment exposure. (D) IC_50_ EA vs AA (t-test *P* = 0.008). n=3 AA vs n=3 EA biological replicates for each experiment, at least 3 independent experiments were performed.

To assess a cell line’s susceptibility/response to bile acid damage, we developed a multivariate score that predicts the amount of bile acid that each cell was exposed to base on its cellular response. This was done using a random forest regression model classifying untreated and 3000 μM BAM treated cells. ROC curve and confusion matrices determined high accuracy of this model (Figure S6A-B). Measurements of compactness and form factor, two top contributing features to the model, demonstrate that cells are elongated post BAM treatment and result in denser CellMask staining (Figure S6C). Predicted response as a percentage of 3000 μM BAM treated cells exhibited dose dependency to actual treatment concentration, which was used to determine IC_50_, representing a concentration in which the majority of cells turnover from healthy-like to damaged-like (see Methods).

To interrogate the response of AA (n=3) or EA (n=3) cells to bile acid, we performed a blinded dose-response (0-3000μM) gradient and analyzed over a million cells per biological specimen. Samples were unblinded only after all data analysis was completed for individual samples. After unblinding samples, we observed that the average IC_50_ for EA derived esophageal cells is 130.4 μM, which is significantly lower than the average IC_50_ observed in AA cell lines (1551 μM) (Figure 4C-D)(*P* = 0.008). These suggest that patient derived stem cells from EA are more susceptible to BAM induced injury than AA stem cells. Even further, it suggests that this differential response mimics clinical observations that AA esophagus are less susceptible to GERD related inflammation and BE when compared to EA.

## Discussion

*In vitro* models of the esophagus have been previously established (Mou *et al*., 2016; Yamamoto *et al*., 2016), however full characterization of the *in vivo* adult human esophagus with *in vitro* derived esophageal cell populations to determine how well cellular heterogeneity is modeled and maintained *in vitro* was unknown. In addition, racial disparities have been described for esophageal disease such as EAC (Spechler *et al*., 2002; Pickens and Orringer, 2003; Abrams *et al*., 2008; Thrift and El-Serag, 2016), but tools to help understand the mechanistic basis for these observations have been lacking. Here, scRNA-seq based characterization of the adult human esophagus *in vivo* and *in vitro* allowed us to characterize the zonation of the stratified epithelium (basal, suprabasal, luminal) and to identify several basal-cell specific markers of the stem cell zone, that had not been previously described (i.e. CAV1, CAV2, COL17A1). Of note, we find that the transcription factor TP63, which is known for its role in basal cell regulation in several tissues such as the skin, lungs and esophagus in mice is not restricted to the basal zone in the human adult esophagus but is more broadly expressed throughout the basal and suprabasal cell zones highlighting species-specific differences in gene/protein expression.

Previous studies describing the growth of human esophagus *in vitro* have relied on the use of canonical markers such as TP63 for validating cell types found in culture. Our single cell analysis suggests that determining the heterogeneity within the *in vitro* environment by using TP63 or by using methods that interrogate the ensemble of mRNA within a population of esophageal cells may be difficult, given that there are many shared molecular and cellular markers within the basal and suprabasal zone. Single cell analysis helped to resolve the different domains within the human esophagus and demonstrated that the expression profile of epithelial cells maintained *in vitro* resemble the basal, suprabasal and proliferative cells of the *in vivo* adult human esophagus.

Clinical studies have shown that AA populations have significantly less esophageal disease including Barrett’s Esophagus and a lower incidence of EAC when compared to EA patients (El-Serag, HB and Sonnenberg, 1997; El-Serag *et al*., 2014). Messenger RNA analyses of cohorts of AA and EA patient-derived esophageal biopsies in healthy and diseased states revealed differences in the mRNA expression of the enzyme GSTT2 in these patients (Ferrer-Torres *et al*., 2019), with the AA biopsies possessing significantly higher GSTT2 mRNA levels. However, the cellular tools to study the mechanisms underlying these racial differences and to interrogate GSTT2 in both AA and EA tissue models, were lacking. This is primarily due to a lack of stem-cell biobanks that represent the diversity of the human population. To address this, we sought to establish a comprehensive and diverse stem-cell biobank of esophageal samples. To this end, we developed a biobank of esophageal cell lines from across the diversity of the human population, and from a wide range of healthy and diseased states (Table 1).

To further leverage this biobank, we developed and implemented a high-content bile-acid injury response assay *in vitro*. Importantly, the *in vitro* findings that AA-derived stem cells are significantly less sensitive to bile-acid induced injury than EA-derived stem cells reflect clinical observations that AA have lower risk of BE and EAC relative to EA, even when risk factors such as acid reflux are identical (El-Serag *et al*., 2004; Xie *et al*., 2017). *In vitro* cellular models of the human esophagus can therefore be used to assess the effects of damaging and carcinogenic agents in patient derived cells across the spectrum of diversity in the human population. Further studies are needed however, to fully understand and characterize the effects observed across cell lines in this model system. Studies presented here suggest that we can use patient-derived stem cells, and high-throughput models to better understand the racial disparities in disease that have been reported for decades.

## Limitations of the study

During the course of this study, the global SARS-CoV-2 pandemic resulted in a research shut down for several months, and a complete halt to research involving human patients for the majority of the past year. This directly affected the ability for the study authors to collect new specimens over the course of the past year in order to complete new experiments. An important caveat to our study is that the racial identity given to a patient sample (and therefore each cells line), is based on the self-identification of race by individual tissue donors. Therefore, the specific genetic ancestry or ancestries of the samples it is yet to be determined. Lastly, we note that while the biorepository reported herein is diverse, representing individuals from multiple ancestries, it is still dominated by EA samples, and as such, there is stillroom for improvement. Given the relatively high proportion of EA patients seen by the University of Michigan, this may be most easily overcome by partnering with institutions or research groups that have greater diversity in their patient populations.

## Supporting information

Supplemental Table 1. Gene signature of Day 0 clusters

Supplemental Table 2. Epithelial-Extracted gene signature of Day 0 clusters

Supplemental Table 3. 2D in-vitro gene signatures clusters

## Acknowledgements

JRS was supported by institutional funds and by a funding from the Chan Zuckerberg Initiative Seed Network, and by the University of Michigan Center for Gastrointestinal Research (UMCGR) (NIDDK 5P30DK034933). DFT was supported by the Center for Plasticity and Organ Design (CPOD) T32 (T32HD007505) and the Michigan Institute for Clinical & Health Research UL1TR002240; KL2TR002241; and TL1TR002242. The content is solely the responsibility of the authors and does not necessarily represent the official views of the National Institutes of Health.

## Author contributions

Project conceptualization: DFT, JRS

Experimental design: DFT, JRS

Experiments and data collection: DFT, JW, CJZ, MAH, MD, AW, EMH, KK, CLM, MSB, SH, YT, DKT, JL, PDH, JS, JRS

Data analysis and interpretation: DFT, JW, CJZ, JS, JRS

Writing manuscript: DFT, CJZ, JRS

Editing manuscript: all authors

## MATERIALS & METHODS

### Sample collection

Histologically normal biopsies of the esophageal squamous epithelium with matched non-dysplastic Barrett’s esophagus (BE)(adjacent) were collected from consenting men and woman who underwent upper endoscopy between 2017 and 2020 at the time of a scheduled BE screening at the University of Michigan Health System. Samples were collected using protocols approved by the University of Michigan institutional review board (IRB). Fresh samples were either used immediately for single cell dissociation followed by scRNA-seq, or were processed for culture; otherwise, biopsies were cryopsreserved and stored at -80 C until use, at which point they could be thawed and processed for generating new culture lines. In order to cryopreserve biopsies, we mince tissue into fine pieces, and freeze using 1ml of HYENAC+ 20% CBS serum and 10% DMSO. Cryovials are place in Mr. Frosty’s at -80 overnight and moved to liquid nitrogen tanks. For culturing cell lines, we found that fresh biopsies could be processed immediately and grown at 100% success rate. We observed that biopsy samples in HYENAC+CBS media can be kept for 24hrs at 4C and can be grown successfully. Even further, samples can be shipped, at which point viable cell lines could still be robustly established. For access to detail protocols see: https://www.umichttml.org/protocols.

### Patient Samples Information

Patients race was self-identified. For white non-Hispanics (W-NH) we used the nomenclature European American (EA), for Black, we use African American (AA). Biological replicates utilized for single-cell RNA sequencing came from the normal squamous biopsies. For the single cell RNA seq studies, patient characteristics are as follows: Patient #1: 45yo, EA, male; Patient #2: 63yo, EA, female, and Patient #3 is a 64yo, EA, female.

### Tissue Processing and Staining

Frozen samples were placed within a mold containing OCT compound and frozen at -80°C. For sectioning, the tissue block was secured onto the sectioning mount within the cryostat using OCT, and 10 μm thick sections were cut at -20°C and placed onto room temperature superfrost plus microscope slides. These slides were stained in hematoxylin for 1 minute followed by a wash in cold running water for 5 minutes and distilled H2O for an additional 2 minutes. Slides were then dipped in 90% EtOH 10 times and counterstained with 1X Eosin Y for 30 seconds. The tissue was dehydrated in 70% EtOH for 3 minutes x 2 changes, 95% EtOH for 3 minutes x 2 changes, 100% EtOH for 3 minutes x 2 changes, and cleared in Histoclear II for 5 minutes x 2 changes. Cover slides were secured using permount mounting media before imaging.

### Tissue Dissociation for Single Cell RNA Sequencing

To dissociate patient biopsies for single cell RNA sequencing, tissue was placed in a petri dish with ice-cold 1X HBSS (with Mg2+, Ca2+). To prevent adhesion of cells, all tubes and pipette tips were pre-washed with 1% BSA in 1X HBSS. The tissue was minced manually using spring-squeeze scissors before being transferred to a 15mL conical containing 1% BSA in HBSS. Tubes were spun down at 500G for 5 minutes at 10°C, after which excess HBSS was aspirated. Mix 1 from the Neural Tissue Dissociation Kit (Miltenyi, 130-092-628) containing dissociation enzymes and reagents was added and incubated at 10°C for 15 min. Mix 2 from the Neural Tissue Dissociation Kit was added, and the suspension was fluxed through P1000 pipette tips, interspersed by 10 min incubations at 10°C. Flux steps were repeated as needed until cell clumps were no longer visible under a stereo microscope. Cells were filtered through a 1% BSA coated 70μm filter using 1X HBSS, spun down at 500g for 5 minutes at 10°C, and resuspended in 500μl 1X HBSS (with Mg2+, Ca2+). 1mL of RBC Lysis Buffer (Roche, 11814389001) was added and tubes were incubated on a rocker at 4°C for 15 minutes. Cells were spun down at 500G for 5 minutes at 10°C, then washed twice in 2mL 1% BSA, being spun down at 500G for 5 minutes at 10°C each time. A hemocytometer was used to count cells, which were then spun down and resuspended to reach a concentration of 1000 cells/μL and kept on ice.

### Single-cell library preparation

The 10x Chromium at the University of Michigan Advanced Genomics Core facility was then used to create single cell droplets with a target of capturing 5,000-10,000 cells. The Chromium Next GEM Single Cell 3’ Library Construction Kit v3.1 prepared single cell libraries according to manufacturer instructions.

### Sequencing Data Processing and Cluster Identification

The University of Michigan Advanced Genomic Core Illumina Novaseq performed all single cell RNA sequencing. Gene expression matrices were constructed from raw data by the 10x Genomic Ranger with human reference genome (hg19). The Single Cell Analysis for Python was utilized for analysis as previously described by (Wolf, Angerer and Theis, 2017). Filtering parameters for gene count range, unique molecular identifier (UMI) counts, and mitochondrial transcript fraction were implemented for each data set to verify high quality input data. For the remainder of processing, all tissue data sets were combined after organ-specific quality filtering had been performed. Highly variable genes were removed, gene expression levels were log normalized, and effects of UMI count and Mitochondrial transcript function variations were regressed out via linear regression. Z-transformation was then performed on gene expression values before samples were again separated by organ for downstream analysis. The UMAP algorithm (Becht et al., 2019)(McInnes et al., 2018) was utilized alongside Louvain algorithm cluster identification within Scanpy with a resolution of 0.6 (Blondel et al., 2008) to perform a graph-based clustering of the top 10-11 principal components. A full detailed protocol for tissue dissociation for single-cell RNA seq can be found at www.jasonspencelab.com/protocols.

### Tissue preparation, Immunohistochemistry, and Imaging Paraffin sectioning and staining

Patient biopsy/tissues were fixed in 4% paraformaldehyde (Sigma) overnight, washed with PBS, and then dehydrated in an alcohol series: 30 minutes each in 25%, 50%, 75% methanol:PBS/0.05% Tween-20, followed by 100% methanol, 100% ethanol and 70% ethanol. Tissue was processed into paraffin using an automated tissue processor (Leica ASP300). Paraffin blocks were sectioned 7 uM thick, and immunohistochemical staining was performed as previously described (Spence et al., 2009). Briefly, slides were rehydrated in a series of HistoClear, 100% ethanol, 95% ethanol, 70% ethanol, 30% ethanol, DI H2O with 2 changes of 3 minutes each. Antigen retrieval as performed in 1X sodium citrate buffer in a vegetable steamer for 40 minutes. Following antigen retrieval, slides were washed in PBS and permeabilized for 10 minutes in 0.1% TritonX-100 in 1xPBS, blocked for 45 minutes in 0.1% Tween-20, 5% normal donkey serum in 1XPBS. Antibodies used in this study can be found in the Key Resources Table. Primary antibodies were diluted in block and applied overnight at 4°C. Slides were then washed 3 times in 1X PBS. Secondary antibodies and DAPI were diluted in block and applied for 40 minutes at room temperature. Slides were then washed 3 times in 1X PBS and cover slipped with ProLong Gold.

### Hematoxylin and eosin

H&E staining was performed using Harris Modified Hematoxylin (FisherScientific) and Shandon Eosin Y (ThermoScientific) according to manufacturer’s instructions. Alcian blue/PAS staining was performed using the Newcomer supply Alcian Blue/PAS Stain kit (Newcomer Supply, Inc.) according to manufacturer’s instructions. Trichrome staining was performed by the University of Michigan in vivo Animal Core.

### Imaging and image processing (Figure 1-2)

Fluorescently-stained slides were imaged on a Nikon A-1 confocal microscope. Brightness and contrast adjustments were carried out using ImageJ (National Institute of Health, USA) and adjustments were made uniformly across images.

### Schematics and Diagrams

Schematic and diagrams in Figure 4A was modified from BioRender (2021) (Hynds et al., 2018). Supplemental Figure 2 schematic made with Biorender, 2021).

### Quantification and Statistical Analysis

Statistical analyses and plots were generated in Prism 8 software (GraphPad). For all statistical tests, a significance value of 0.05 was used. For every analysis, the strength of p values is reported in the figures according the following: p > 0.05, *p ≤ 0.05, **p ≤ 0.01, ***p ≤ 0.001, ****p ≤ 0.0001. Details of statistical tests can be found in the figure legends. With the exception of scRNA-seq, three HT lines were used across experiments with at least 2-3 independent experiments and at least 2-3 technical replicates per experiment.

### Computational analysis of single-cell RNA sequencing data

#### Overview

To visualize distinct cell populations within the single-cell RNA sequencing dataset, we employed the general workflow outlined by the Scanpy Python package (Wolf, Angerer and Theis, 2017). This pipeline includes the following steps: filtering cells for quality control, log normalization of counts per cell, extraction of highly variable genes, regressing out specified variables, scaling, reducing dimensionality with principal component analysis (PCA) and uniform manifold approximation and projection (UMAP) (McInnes et al., 2018), and clustering by the Louvain algorithm (Blondel et al., 2008).

#### Sequencing data and processing FASTQ reads into gene expression matrices

All single-cell RNA sequencing was performed at the University of Michigan Advanced Genomics Core with an Illumina Novaseq 6000. The 10x Genomics Cell Ranger pipeline was used to process raw Illumina base calls (BCLs) into gene expression matrices. BCL files were demultiplexed to trim adaptor sequences and unique molecular identifiers (UMIs) from reads. Each sample was then aligned to the human reference genome (hg19) to create a filtered feature bar code matrix that contains only the detectable genes for each sample.

#### Quality control

To ensure quality of the data, all samples were filtered to remove cells expressing too few or too many genes (Figure S1/Figure 1/Figure S2/Figure2/Figure S3/Figure 3/FigureS4: <500, >7500, or a fraction of mitochondrial genes greater than 0.2.

#### Normalization and Scaling

Data matrix read counts per cell were log normalized, and highly variable genes were extracted. Using Scanpy’s simple linear regression functionality, the effects of total reads per cell and mitochondrial transcript fraction were removed. The output was then scaled by a z-transformation. Following these steps, a total of (Figure S1—9039 cells, 3897 genes; Figure 1/Figure S2 (extracted)—7796 cells, 2651 genes; Figure 2/Figure S3A-C—10550 cells, 4269 genes; Figure S3E-F—HT239(4617cells, 4845 genes), HT344(895 cells, 3034 genes), HT328(4133 cells, 5195 genes), Figure 3C-H—7389 cells, 3413 genes, FigureS4—19589 cells, 3486 genes.

#### Variable Gene Selection

Highly variable genes were selected by splitting genes into 20 equal-width bins based on log normalized mean expression. Normalized variance-to-mean dispersion values were calculated for each bin. Genes with log normalized mean expression levels between 0.125 and 3 and normalized dispersion values above 0.5 were considered highly variable and extracted for downstream analysis.

#### Batch Correction

We have noticed batch effects when clustering data due to technical artifacts such as timing of data acquisition or differences in dissociation protocol. To mitigate these effects, we used the Python package BBKNN (batch balanced k nearest neighbors)(Polański et al., 2020). BBKNN was selected over other batch correction algorithms due to its compatibility with Scanpy and optimal scaling with large datasets. This tool was used in place of Scanpy’s nearest neighbor embedding functionality. BBKNN uses a modified procedure to the k nearest neighbors’ algorithm by first splitting the dataset into batches defined by technical artifacts. For each cell, the nearest neighbors are then computed independently per batch rather than finding the nearest neighbors for each cell in the entire dataset. This helps to form connections between similar cells in different batches without altering the PCA space. After completion of batch correction, cell clustering should no longer be driven by technical artifacts.

#### Dimension Reduction and Clustering

Principal component analysis (PCA) was conducted on the filtered expression matrix followed. Using the top principal components, a neighborhood graph was calculated for the nearest neighbors Figure S1– 16 principal components, 30 neighbors; Figure 1/ Figure S2—9 principal components, 11 neighbors; Figure 2/ Figure S3—30 principal components, 16 neighbors; Figure 3C-H—11 principal components, 15 neighbors; Figure S4—16 principal components, 30 neighbors. BBKNN was implemented when necessary and calculated using the top 50 principal components with 3 neighbors per batch. The UMAP algorithm was then applied for visualization on 2 dimensions. Using the Louvain algorithm, clusters were identified with a resolution of (Figure S1—0.3; Figure 1/Figure S2—0.2; Figure 2/Figure S3—0.3; Figure 3—0.4, and Figure S4—0.3)

#### Cluster Annotation

Using canonically expressed gene markers, each cluster’s general cell identity was annotated. Markers utilized include epithelium (CDH1), mesenchyme (VIM), neuronal (POSTN, S100B, STMN2, ELAV4), endothelial (ESAM, CDH5, CD34, KDR), and immune (CD53, VAMP8, CD48, ITGB2).

#### Sub-clustering

After annotating clusters within the UMAP embedding, specific clusters of interest were identified for further sub-clustering and analysis. The corresponding cells were extracted from the original filtered but unnormalized data matrix to include (Figure 1A/S2 – 9039 cells, 3897 genes). The extracted cell matrix then underwent log normalization, variable gene extraction, linear regression, z transformation, and dimension reduction to obtain a 2-dimensional UMAP embedding for visualization.

### High-Content Imaging

ECAD quantifications (Figure 2G-H) Cells were cultured, fixed and stained with Hoechst 33342, Cell Mask Deep Red, and ECAD antibody + secondary (488) in PerkinElmer CellCarrier-384 Ultra Microplates (6057300). Automated imaging was done using ThermoFisher Scientific CellInsight CX5 High-Content Screening Platform. The open-source CellProfiler (3.1.9) was used for cell object segmentation and measurements of ECAD intensity. Threshold value for ECAD positivity was determined based on measured values for visually positive cells in a subset of images. Cells with a measured ECAD value higher than the threshold was determined to be positive.

### Bile-Acid Treatment

(Figure 4B-D, Figure S5)

Cells were cultured at an initial seeding density of 2,000 cells/well on PerkinElmer CellCarrier-384 Ultra Microplates and grown for 48 hrs to 80% confluence. Cells were then treated with a 10-point, 1:3 serial dilution, dose range of a bile-acid mixture from 0 to 3000 μM with 4 well-level replicated. After 48 hrs of treatment, cells were fixed and stained with Hoechst 33342, Cell Mask Orange (nuclei and cell boundary segmentation), and NFkB (cell stress). Confocal images were analyzed with Cell Profiler

3.1.9 to obtain nuclei and cell morphological features. Cells were then grouped by treatment and analyzed using linear discriminant analysis in JMP Pro 14 which report a percent misclassification of cells, i.e. the percentages of cells wrongly attributed to the wrong treatment group. Lower misclassifications are representative of more distinctive morphological features, and therefore, higher prediction accuracy by any treatment described a greater morphological perturbation from treatment. Figure S5; To determine the percentage accuracy, each cell was classified as exhibiting an untreated or high bile-acid treated phenotype based on linear discriminant analysis in JMP. Prediction accuracies were determined based on the inverse of the reported misclassification rate. Higher accuracies represent a more distinctive phenotype between untreated vs treated cell groups.

CellProfiler was used for automated nuclei and cell segmentation followed by measurements of morphological features (size and shape), intensity and distribution of stains amassing 1,400 unique measured features per-cell. These measured features were normalized and filtered for low-variance and high-correlation with Konstanz Information Miner (KNIME)(Berthold et al., 2006). Random forest models were designed based on a regression model of untreated cells vs 3000 μM treated cells, and individual cells were scored along this model for a percent response between 0 and 3000 μM treated cells. A 80/20 split was done where 20% of the data was used in model creation and 80% for model validation. ROC curve and confusion matrix both show high model accuracy (Figure S6). Per-well averages of predicted percentage response were obtained and plotted vs actual treatment value in GraphPad Prism 8. IC50 values were calculated using GraphPad’s nonlinear regression curve fitting.

## Supplemental Figures

**Supplemental Figure 1. Human esophageal biopsies characterized by scRNA-seq**. (A) Patient biopsies (n=2) were sequenced on the day of biopsy (day 0), a total of 9,039 cells included in the analysis after filtering, with 3,897 genes expressed per cell. Louvain clustering predicts seven molecularly distinct clusters (each cluster was determined using the top 200 genes differentially expressed in each cluster (P<0.01)). (B) Individual patient samples have similar distribution of cells to each cluster. <u>*x*</u> denotes the average contribution of both samples to the clusters. (C-D) Identification of epithelial cells (CDH1+) and lamina propria/mesenchymal cell types (VIM+). (E-F) Dot plots of the top 5 genes expressed in each cluster (E) and feature plots of selected marker genes (F) were used to annotate each cluster (Table S1) (G) Protein expression of top markers identified using scRNA-seq for the different epithelial zones of the esophagus (data from the Human Protein Atlas)(Uhlen, 2005; Uhlen *et al*., 2015). Scale bar represents 50μm (20X)

**Supplemental Figure 2. Characterizing the adult human esophagus epithelium**. (A) Feature plots for scRNA-seq related to Figure 1 showing molecular markers enriched in epithelial clusters: Basal cells (Cluster 2 – CAV2-enriched) a suprabasal proliferative zone (Cluster 1 - LY6D+), a middle-zone (Cluster 0 - KRT4+) and a completely differentiated luminal zone (Cluster 3). Note, TP63 is expressed throughout Clusters 1, 2 and 3, and is not specific to the basal cells. (C) Validation using immunofluorescence to identify the different zones of the human adult esophagus. CAV1, CAV2, and COL17A1, mark basal stem cells of the esophagus. The LY6D zone starts just one cell-layer above the basal cells, denoting the early-suprabasal layer. KRT4 is expressed in the suprabasal/transitional layer, and CRNN is expressed in the completely differentiated luminal zone of the esophagus. ECAD is used to identify the epithelial cells, DAPI for nuclei, and TP63 is expressed broadly within the esophagus. (D) Summary schematic of different epithelial zones of the esophagus with their corresponding markers identified by scRNA-seq and validated by immunofluescence. Scale bars in 100 (top), 50 (middle) and 30 (lower) µm, respectively (staining for each marker combination was validated in n=3 biological replicates).

**Supplemental Figure 3. *In vitro* cells are analogous to their *in vivo* counterparts and are enriched for basal cells**. (A) Dot plots of the top 5 genes expressed in each cluster of cells *in vitro*. (B) Louvain clustering and UMAP visualization of predicts cell clustering for *in vitro* grown samples (n=3). (C) Feature plots of genes associated with basal stem cells including *KRT15, TP63, CAV2*, and *ITGB4*. (D) Validation of protein expression *in vitro* for CAV2, TP63 (yellow) and co-stained for ECAD (red) and DAPI (grey). Scale bars represent 2mm for CAV2, 100µm for KRT5 and 200µm for TP63 images. (E) Clustering of individual *in vitro* patient samples with feature plots for CDH1 and VIM expression levels and percentage of cells expressing each marker. (F) Feature plots for scRNA-seq data for individual patient cell lines, for expression of basal cell and epithelial markers (*COL17A1, KRT14, TP63, KRT15*) of the esophagus per patient in-vitro sample (G) Protein expression patterns using Human Protein Atlas immunohistochemistry (IHC) for COL17A1, KRT14, TP63, KRT15 in adult in the *in vivo* human esophagus from the Human Protein Atlas (Uhlen, 2005; Uhlén *et al*., 2015) demonstrating expression patterns of markers identified by IHC.

**Supplemental Figure 4. (**A) All epithelial cells were extracted from *in vivo* (day 0) cells and from cultured *in vitro* cells and were batch corrected using BBKNN. (B) Louvain clustering and UMAP visualization revealed 4 predicted clusters. Molecular characterization (See panels C, D) identifies Cluster 1 as expressing genes at the basal zone, Cluster 3 expressing proliferative genes, Cluster 0 expressing genes of the suprabasal cells and Cluster 2 expressing genes of luminal cells. (C-D) Top 5 enriched genes expressed in each cluster with feature plots (D) for selected cluster-associated genes, including CAV1, KI67, LY6D, KRT4, and CRNN. (E) Quantification of number of cells contributing to each cluster from *in vivo* or *in vitro* samples. There was a significant enrichment for the proportion of basal cells in Cluster 1 from in the *in vitro* cells compared to *in vivo* cells (4.52% to 37.35, respectively). n=2 *in vivo* and n=3 *in vitro* biological replicates.

**Supplemental Figure 5. Bile-acids treatment of patient derived esophagus cell cultures**. A) Prediction accuracies (see methods) of measured morphological features of untreated vs 300⌈M bile-acid mixture treated esophageal cells. Cells treated for 48 hrs with 300⌈M of individual bile-acids, cumene-hydroperoxide, a bile-acid mixture encompassing all other bile-acids, and a bile-acid/cumene hydroperoxide mixture were treated, fixed and stained with Hoechst 33342 and HCS Cell Mask Orange. B) Representative images of cells stained with Cell Mask Orange after no treatment (untreated control) and treated with 300⌈M bile-acid mixture, demonstrating a typical morphological perturbation by treatment. (C) Representative images are shown of individual esophageal epithelial cells (n = 3 for EA and AA) 48hrs after treatment with 0, 300, and 3000⌈M of bile-acid mixture. Images are stained by Hoechst 33342 (Nuclei, cyan) and HCS Cell Mask Orange (magenta) respectively. Images are obtained using a Yokogawa CQ1 Benchtop High-content Analysis system and are maximum projections of 10 highest intensity Z-stacks across a 20 ⌈m range. n=3 AA vs n=3 EA biological replicates for each experiment, at least 3 independent experiments were performed.

**Supplemental Figure 6. Random forest model accuracy and parameters**. A) ROC curve of random forest model used to classify percent bile acid response on a per cell level with an AUC of 0.925. B) Confusion matrix comparing the number of accurate and misclassified cells. C) Measurements of four of the most prominent features used by the model to classify treated and untreated cells are shown across all cells. Wilcoxon-Mann-Whitney Test on well-level averages was done to determine significance between untreated and treated as well as between cell line ethnic origin (* = p < 0.05, ** = p < 0.01). Form factor and compactness (top), both measurements of the cell’s shape, shift dramatically with BAM treatment but non-discriminant across cell lines. Features in relation to Cell Mask Orange staining (bottom) also result in significant changes with BAM treatment but perturbations differ between CAU and AA lines resulting in measured significance between the lines only post-treatment.

## Tables

Table 1. Clinical Characteristics of Primary Human Esophageal Samples for 2D or 3D *in-vitro* culture

Supplemental Table 1. Gene signature of Day 0 clusters

Supplemental Table 2. Epithelial-Extracted gene signature of Day 0 clusters

Supplemental Table 3. 2D in-vitro gene signatures clusters

