## Supplemental Figure 1-6 for "*In vitro* models of the human esophagus reveal ancestrally diverse response to injury"

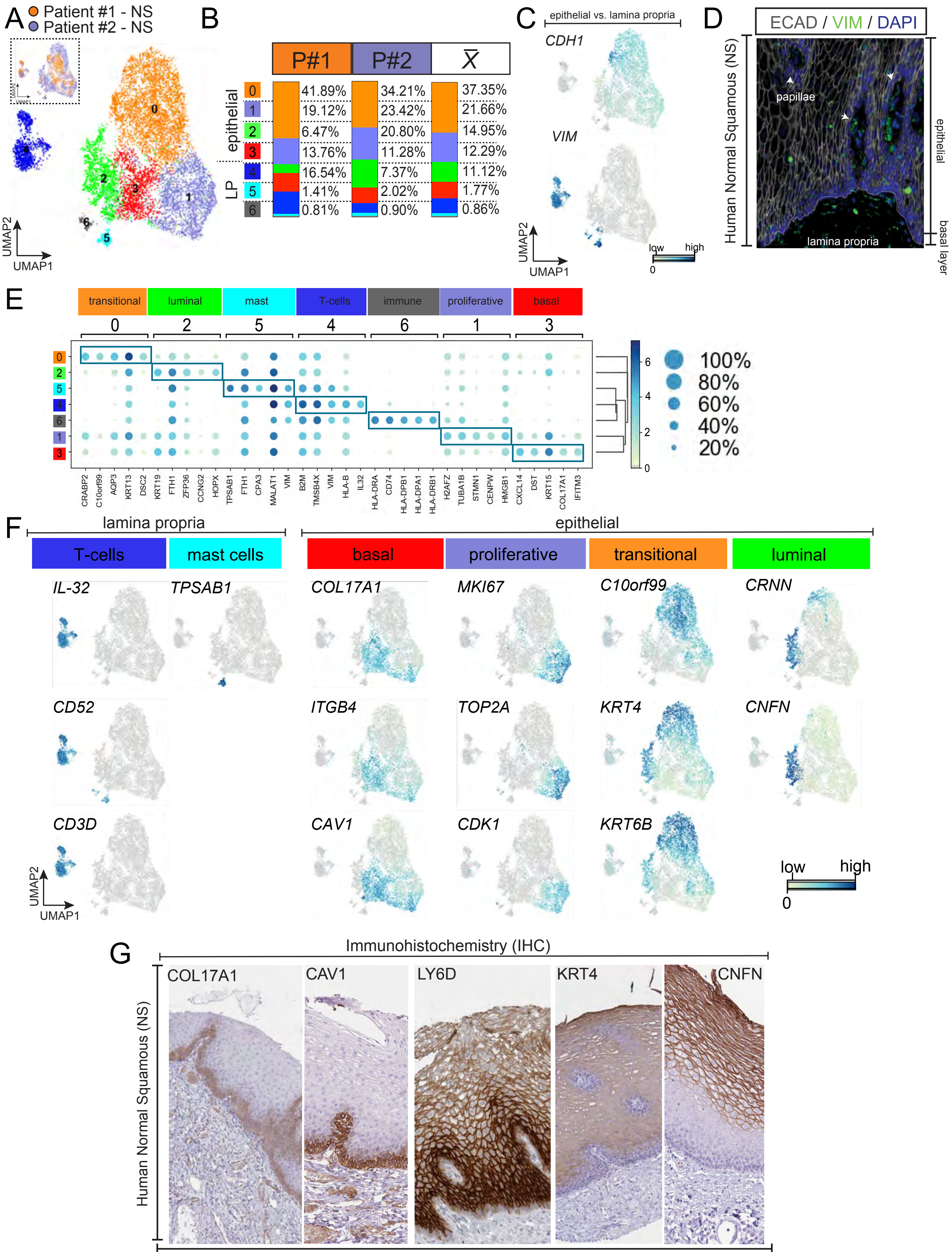


Supplemental Figure 1

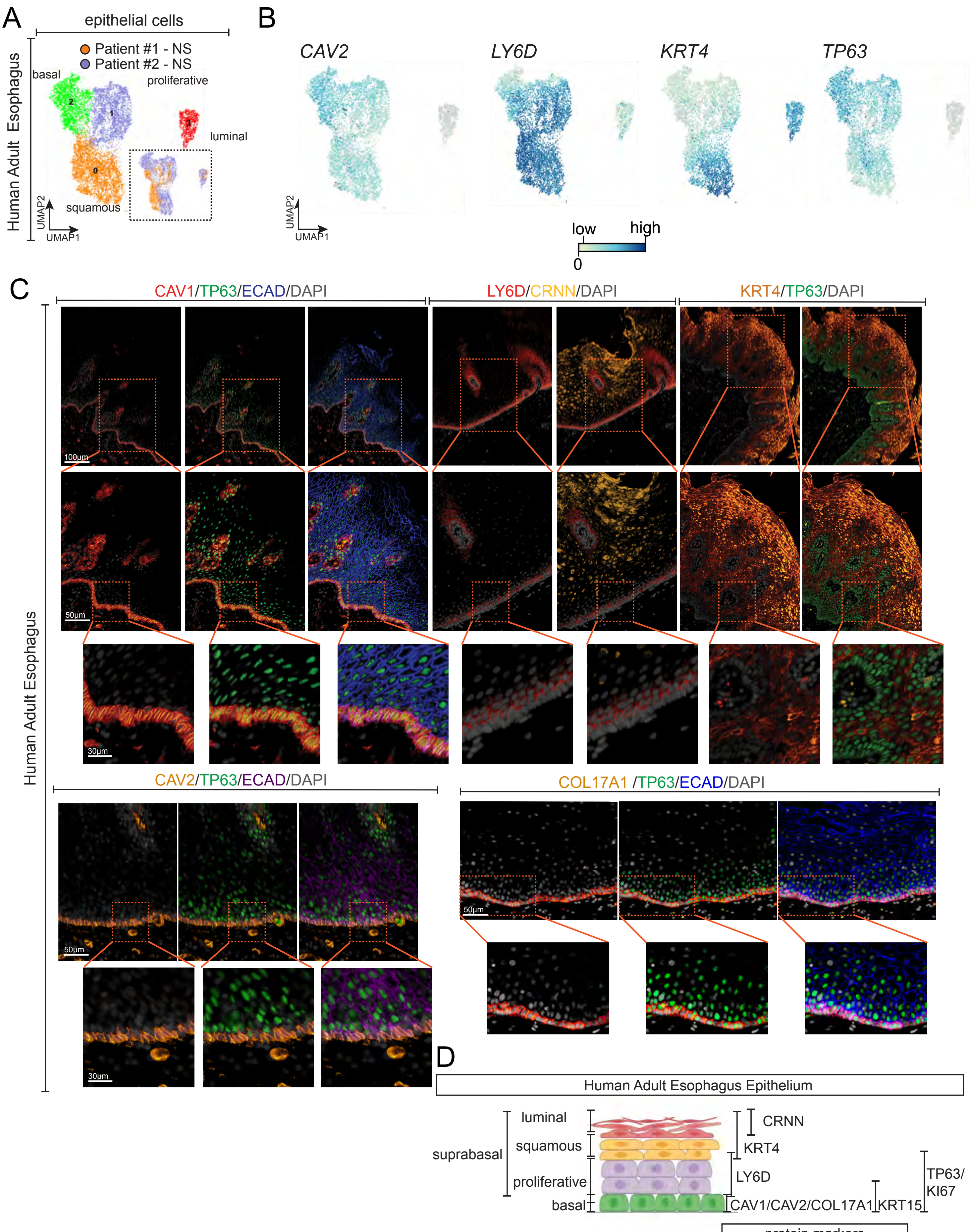


Supplemental Figure 2

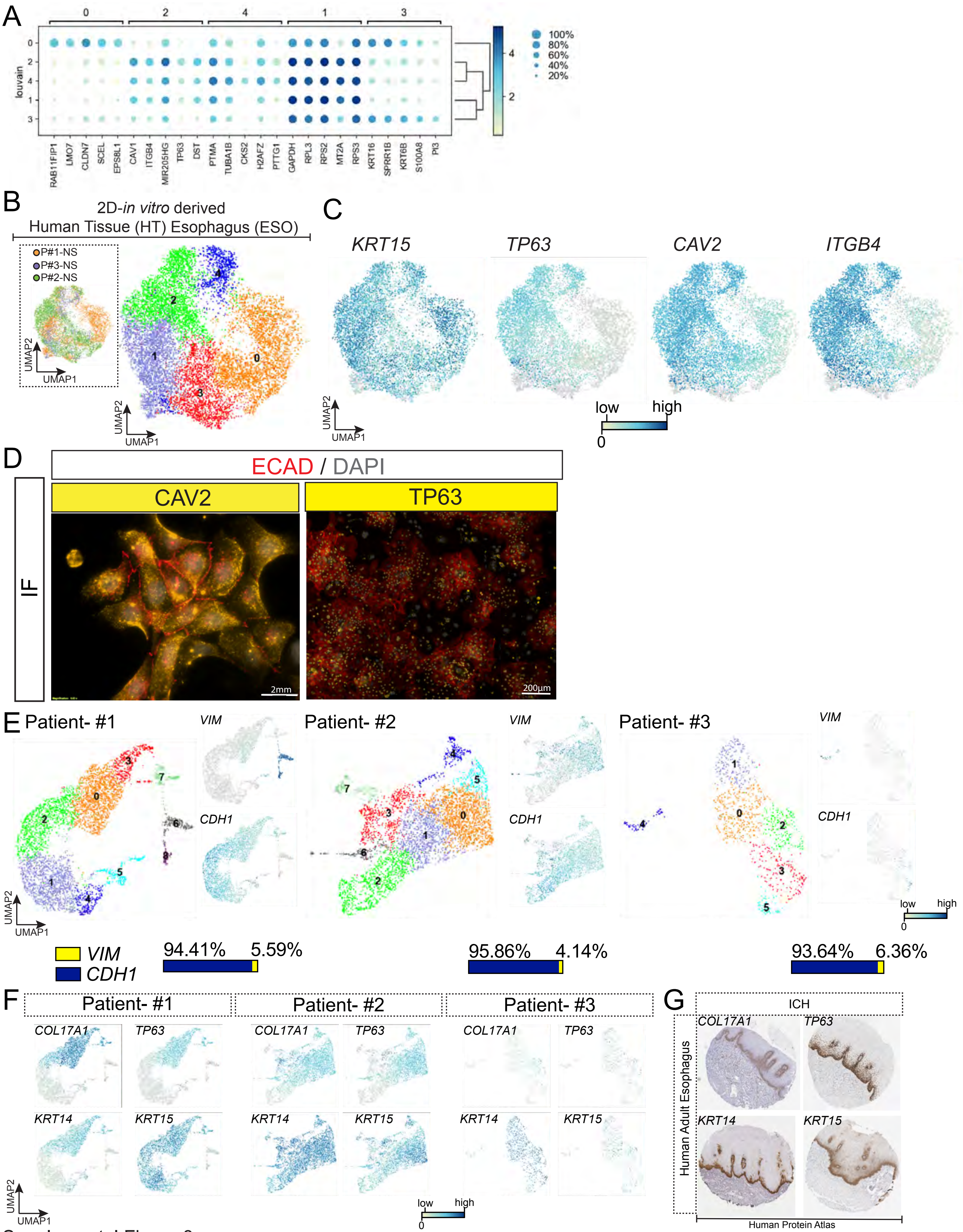

Supplemental Figure 3

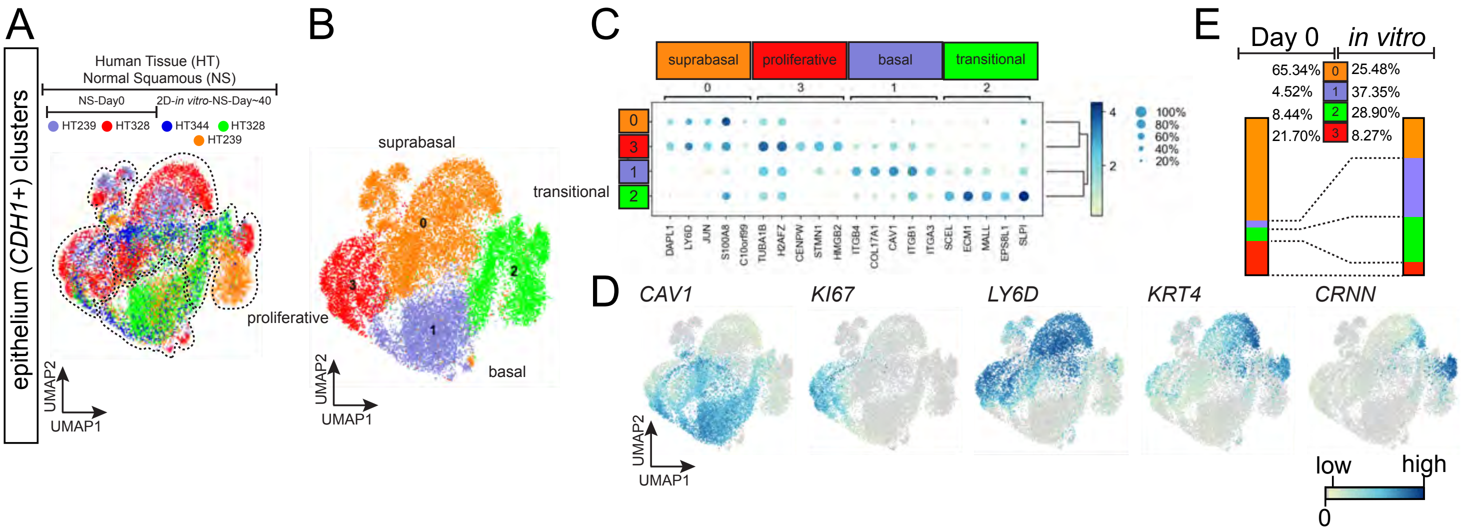


Supplemental Figure 4

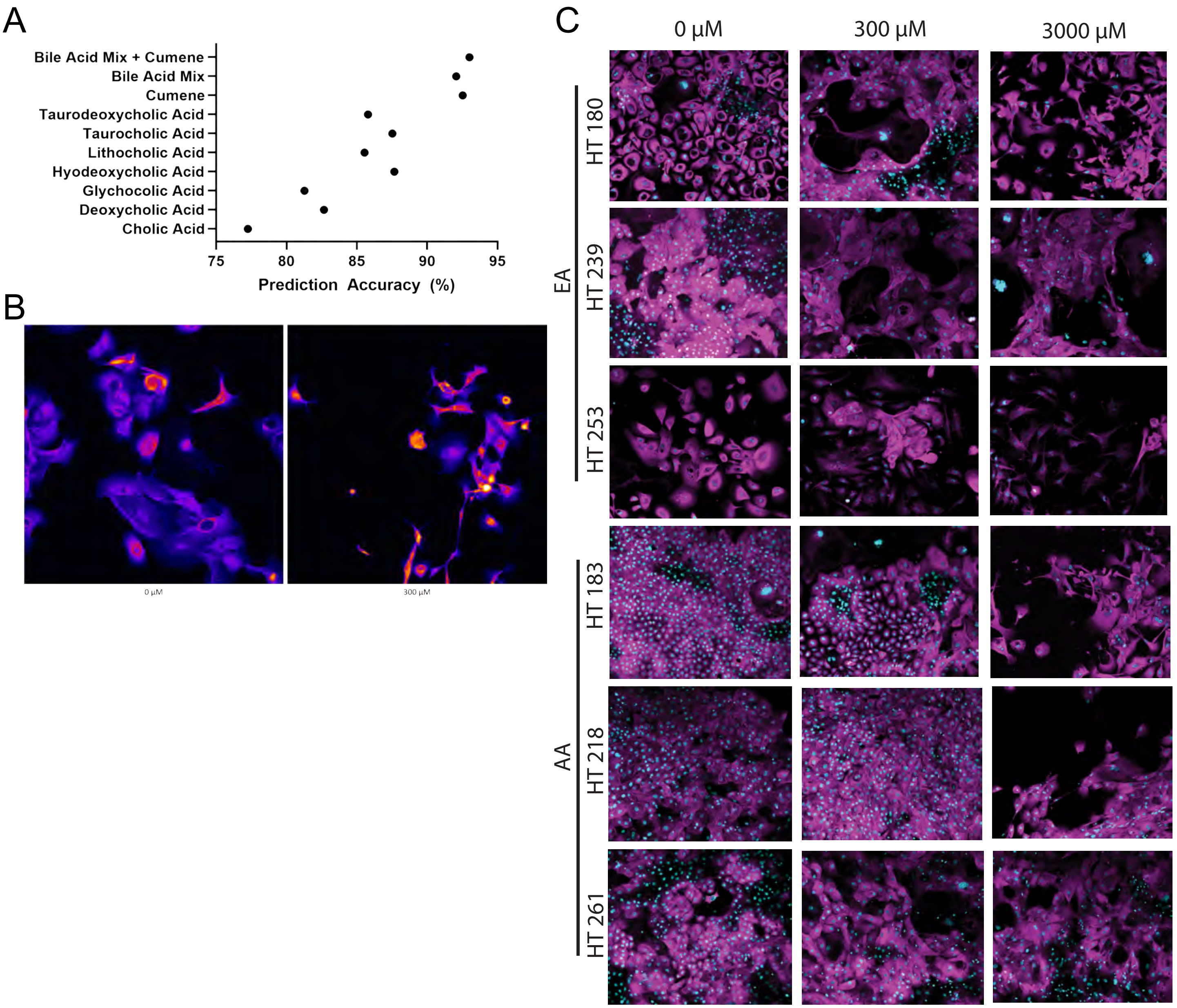


Supplemental Figure 5

A

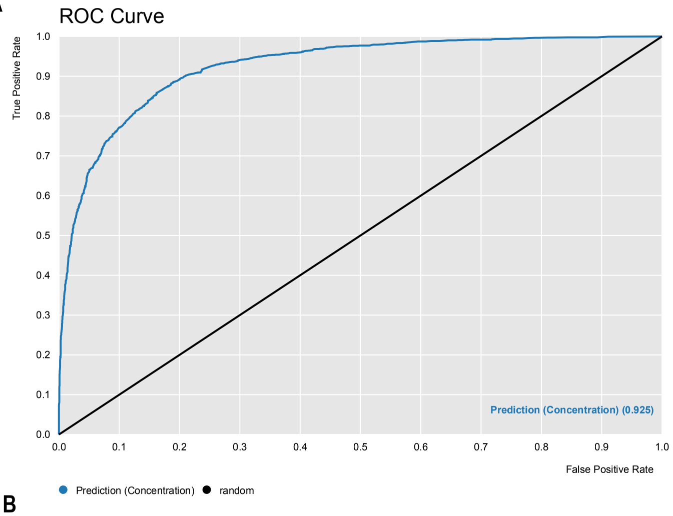


B

| | Vehicle (Predicted) | BAM (Predicted) |
|---|---|---|
| **Vehicle (True)** | 62558 | 7375 |
| **BAM (True)** | 1050 | 30355 |

C

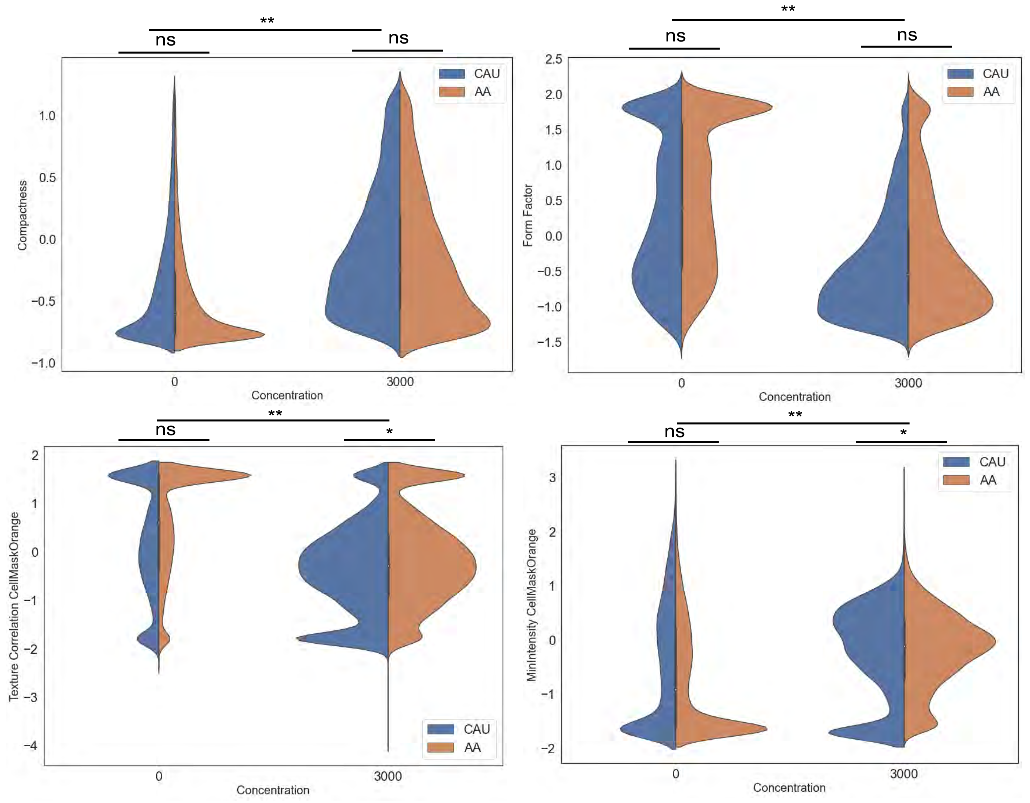
